# Heterochromatin re-organization associated with the transcriptional reprogramming under viral infection in *Arabidopsis*

**DOI:** 10.1101/2023.08.30.555647

**Authors:** Maria Luz Annacondia, Vasti Thamara Juarez-Gonzalez, Jinping Cheng, Juan Luis Reig-Valiente, German Martinez

**Author notes:** Department of Plant and Environmental Sciences, Copenhagen Plant Science Centre, University of Copenhagen, Frederiksberg, Denmark (Maria Luz Annacondia).

## Abstract

Epigenetic mechanisms are key regulators of genomic integrity and genic expression. Emerging evidence shows that epigenetic regulation is an important component of the transcriptional reprogramming during stress. Despite this, the overall stress-induced reprogramming of the different epigenetic marks and their targets are unknown. Here, we uncovered multiple epigenetic changes taking place during viral infection in *Arabidopsis thaliana* and their connection with gene expression. We find that cucumber mosaic virus (CMV) infection induces an overall reorganization of the repressive epigenetic marks H3K9me2, H3K27me3, and DNA methylation, which interact between them and are dynamic during infection. Overall, these epigenetic changes are involved in the reprogramming of the transcriptional program to adapt to the biotic stress, and might ensure genome stability through the transcriptional control of transposable elements (TEs). Mechanistically, we demonstrate that the catalytic component of the Polycomb Repressive Complex 2 (PRC2) CURLY LEAF (CLF) mediates the transcriptional repression of genes gaining H3K27me3 during viral infection and that mutants on that component induce resistance against CMV. Altogether, our results provide a complete picture of the epigenetic changes that occur during biotic stress and exemplify the overall dynamism of epigenetic regulation in eukaryotic organisms.

## Main

Eukaryotic organisms adapt to their environment by modulating their transcriptional program^1,2^. An extreme case of environmental signals are stresses, which usually induce a myriad of development defects due to the elicitation of the defense response^2^. In plants, biotic stresses, in general, and viruses, in particular, induce alterations that result from their hijacking of the host molecular machinery to complete their life cycle^3,4^. To counteract viral infections, plants have several overlapping defense mechanisms that inhibit the accumulation of viral genomes and/or modulate gene expression^5–9^. Due to their dynamic nature, epigenetic mechanisms have been proposed as important regulators of the defense response during stress^10–12^. Nevertheless, how epigenetic mechanisms (especially histone marks) mediate this transcriptional control is poorly understood.

Epigenetic regulation comprises different mechanisms that are fundamental for the maintenance of genome stability^13–15^ and the regulation of the transcriptional program in crucial biological processes^16–21^, including the response against stress^10–12^. Among the different epigenetic mechanisms, the influence of DNA methylation over the stress-induced transcriptional reprogramming has been widely studied in different plant-pathogen interactions such as the ones mediated by bacteria ^22–25^, fungi ^26^, nematodes ^27^, insects ^28^, viroids ^29–31^ and viruses ^32–34^. Both hyper-and hypomethylation of gene regulatory^23,35^ and gene body regions^36^ have been connected to transcriptional changes relevant to the response to stress. In addition, although less studied in plants, histone modifications have also been suggested as important players in the regulation of the defense response, with histone deacetylases playing a role in the transcriptional activation of several stress-responsive genes^37–40^. Histone re-organization during stress is an important part of the stress-induced transcriptional reprogramming in multiple eukaryotic organisms including mammals, Neurospora, and Drosophila, where repressive histone marks located in both facultative and constitutive heterochromatin (mainly the trimethylation of lysine 27 of histone H3, H3K27me3, and the di/trimethylation of lysine 9 of histone H3, H3K9me2/3, respectively) are redistributed under different types of stresses^41–46^. Nevertheless, the dynamics of these two marks under stress or their connection to DNA methylation is unexplored in plants.

Since epigenetic regulation involves the interaction between multiple overlapping regulatory mechanisms, here, to fully understand the role and importance of these mechanisms, we studied the interaction between DNA methylation; the main repressive histone marks H3K9me2 and H3K27me3 and the transcriptional response to cucumber mosaic virus (CMV) infection in *Arabidopsis thaliana*. We discovered that H3K9me2 and H3K27me3 are more dynamic than DNA methylation and exert both a genome-protective role and a gene regulatory effect. In line with this role, we found that the catalytic component of the PRC2 complex CURLY LEAF (CLF) is resistant to CMV infection and a key player in the transcriptional regulation of CMV-responsive genes. Hence, the interaction between different epigenetic mechanisms is a key aspect of stress-responsive transcriptional reprogramming.

## RESULTS

### Different epigenetic pathways control CMV susceptibility in *Arabidopsis thaliana*

Previous studies indicated that DNA methylation could be an important component of the plant viral-induced transcriptional response^47,48^. To understand the contribution of epigenetic regulation to the resistance against viral infection, we analyzed the susceptibility to CMV infection (the highly symptomatic subgroup I strain Fny-CMV^49^, herein referred to as CMV) of mutants covering different epigenetic marks including DNA methylation (*polIV*, *ago4*, *drm2,* and *ddc*), H3K9 methylation (*ddm1, kyp* and *cmt3*) and H3K27 methylation (*clf* and *ref6*) homeostasis (Figs. 1a and b). CMV-infected *Arabidopsis* plants are smaller and more compact than healthy plants^49^, hence, measuring the rosette radius is a good proxy of the viral symptomatology (Fig. 1a and Sup Fig. 1a). Quantification of the susceptibility at two different infection points, 10 days post-infection (dpi, onset of the viral symptomatology) and 20 dpi (advanced developed symptoms) showed marked resistance to CMV infection of some of the mutant lines (Fig. 1b). At 10 dpi, the rosette radius of four different mutants covering the three epigenetic pathways under study (*clf*, *ref6*, *polIV,* and *ddm1*) did not show the expected significant differences between mock and infected plants (Fig. 1b top panel). At a later infection time (20 dpi), only *clf* retained a lack of significant difference between mock and infected plants, indicating a resistance/tolerance to CMV infection (Fig. 1b lower panel). In accordance with an epigenetic component associated with CMV infection, infected plants showed a significant decondensation of H3K9me2 (Sup Fig. 1b). These changes were not attributed to a differential accumulation of CMV in the mutant backgrounds (Sup Fig. 2a and b). Overall, these results suggest that different epigenetic pathways, including DNA methylation, chromatin maintenance and, especially, H3K27 methylation, play an essential (and previously uncharacterized) role in mediating resistance against CMV infection.

**Figure 1.**
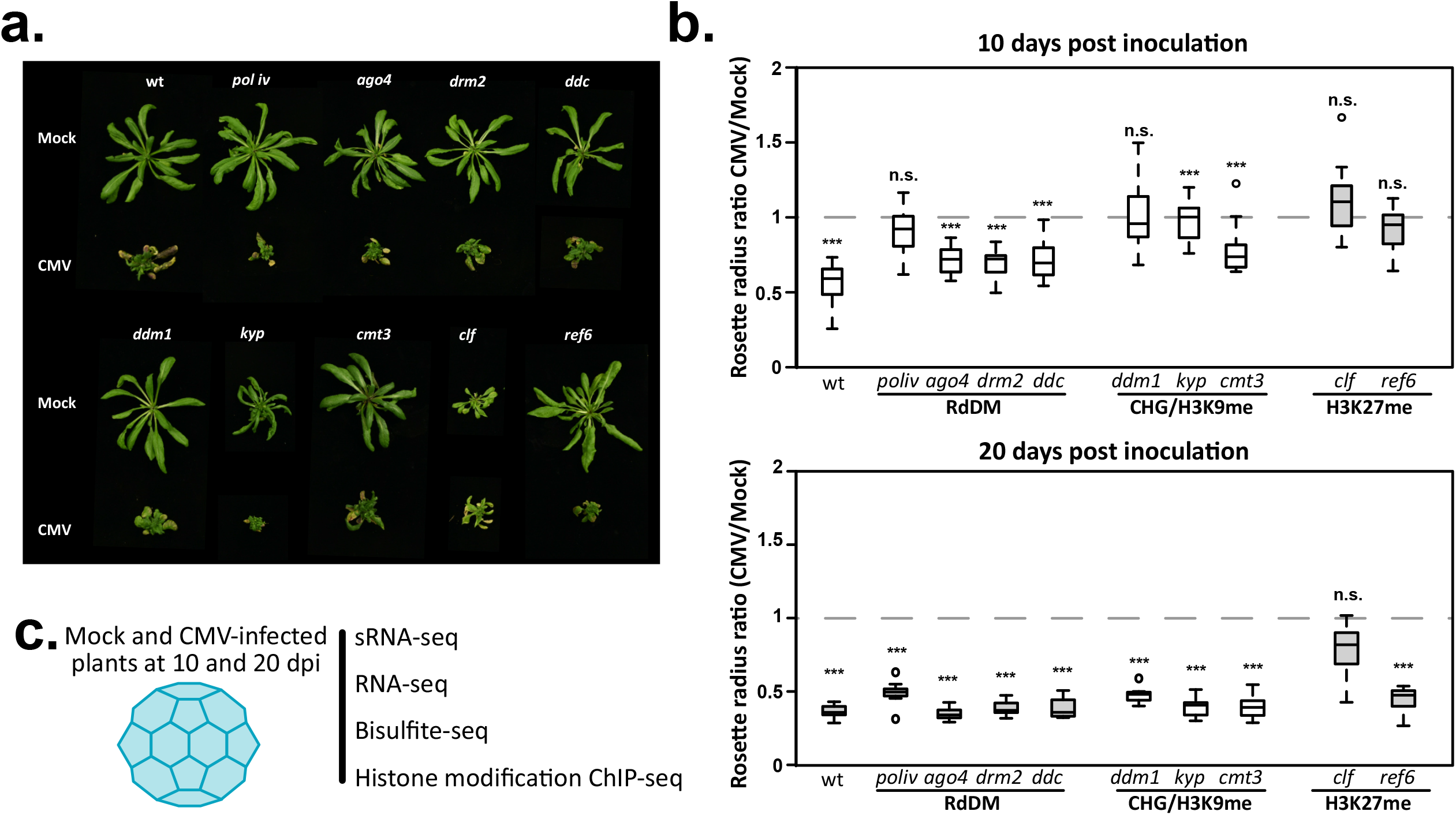
Epigenetic mutants show different susceptibility toward CMV infection. **a.** Representative pictures of the mock and CMV-infected studied genotypes at 20 dpi. **b.** Boxplot of the rosette radius ratio in mock/CMV-infected plants at 10 dpi (top panel) and 20 dpi (bottom panel). Asterisks (***) indicate a p-value <0.005; n.s.=“non-significant”. p-values were calculated using an unpaired t-test. **c.** Experimental setup to explore the overall extent of CMV-induced epigenomic changes. Mock and CMV-infected tissues at 10 and 20 dpi were collected and used to generate whole-genome high-throughput sRNA-, RNA-, bisulfite- and chromatin immunoprecipitation-sequencing.

### Transcriptional reprogramming during CMV infection is characterized by the activation of the defense response

To fully understand how DNA methylation and histone marks could play a role during CMV infection, we used high-throughput sRNA, RNA, whole-genome bisulfite (WGBS), and chromatin immunoprecipitation (ChIP, for the main heterochromatin determinants H3K9me2 and H3K27me3) sequencing from *Arabidopsis thaliana* plants infected with CMV at both 10 and 20 dpi (Fig. 1c).

Previous analysis of the transcriptomic response against CMV infection indicated a strong deregulation of gene expression with multiple molecular functions affected that included metabolic processes, transcription factor binding, hormone signaling and photosynthesis-associated processes^50,51^. Our RNA sequencing analysis (Sup Fig. 3a) indicated that a substantial number of genes were differentially expressed (adjusted p-value <0.05, herein DEGs) under CMV infection: 886 and 723 at 10 and 20 dpi, respectively (Fig. 2a and Sup Table 1). A similar number of genes were up- and downregulated at both 10 (426 and 460) and 20 (309 and 414) dpi, with a considerable overlap of misregulated genes both in the up and downregulated fractions (56% of average overlap between 10 and 20 dpi for up- and downregulated genes, Fig. 2b). Gene ontology (GO) categorization of CMV-responsive genes according to molecular function highlighted a preference of genes involved in metabolic regulation and signaling receptor activity being commonly enriched among upregulated genes, while genes involved in structural molecular activity, translation factor, oxygen and RNA binding are commonly downregulated (Fig. 2c, enriched genes according to their overall genome representation). Our analysis highlighted a series of transcription factors (TFs) that were identified as DEGs (up- or downregulated) and that had either a general activity during infection or specific activity at either 10 or 20 dpi and might be important in the orchestration of the transcriptional response against CMV, such as HAT1 (downregulated at 10 dpi)^52^ or WRKY70 (upregulated at both 10 and 20 dpi)^53^ (Fig 2d). We identified several TFs which were previously uncharacterized as DEGs under CMV infection and that are key for proper plant development, such as SHY2 (downregulated at 10 and 20 dpi) or several WRKY family members (6, 38, 51, and 75, all upregulated at 10 and 20 dpi) (Fig 2d.). Additionally, we identified several genes associated with different epigenetic pathways that experience up- or downregulation under CMV infection, but that were not categorized as DEGs (Sup. Fig 3b). These include the upregulation (>0.5 log2 (fold change)) of the H3K9 demethylase JUMONJI 26 (JMJ26), the importin/exportin HASTY (HST), and the linker histone 1.2 (H1.2) at 10 dpi; the upregulation of DSRNA-BINDING PROTEIN 5 (DRB5), AGO3, DEMETER-LIKE 2 (DML2), HISTONE DEACETYLASE 6 (HDA6) and AGO2 (the only real DEG from this group of genes) at both 10 and 20 dpi; and the upregulation of DRM2 at 20 dpi (Sup Fig. 3b); and the downregulation (>-0.5 log2 (FC)) of the H3K27 demethylase JUMONJI 13 (JMJ13), AGO7, HYPONASTIC LEAVES 1 (HYL1), DDM1 at 10 dpi and of CLF, MET1 and HISTONE-LYSINE N-METHYLTRANSFERASE ATXR2 (ATXR2) at 20 dpi (Sup. Fig 3b).

**Figure 2.**
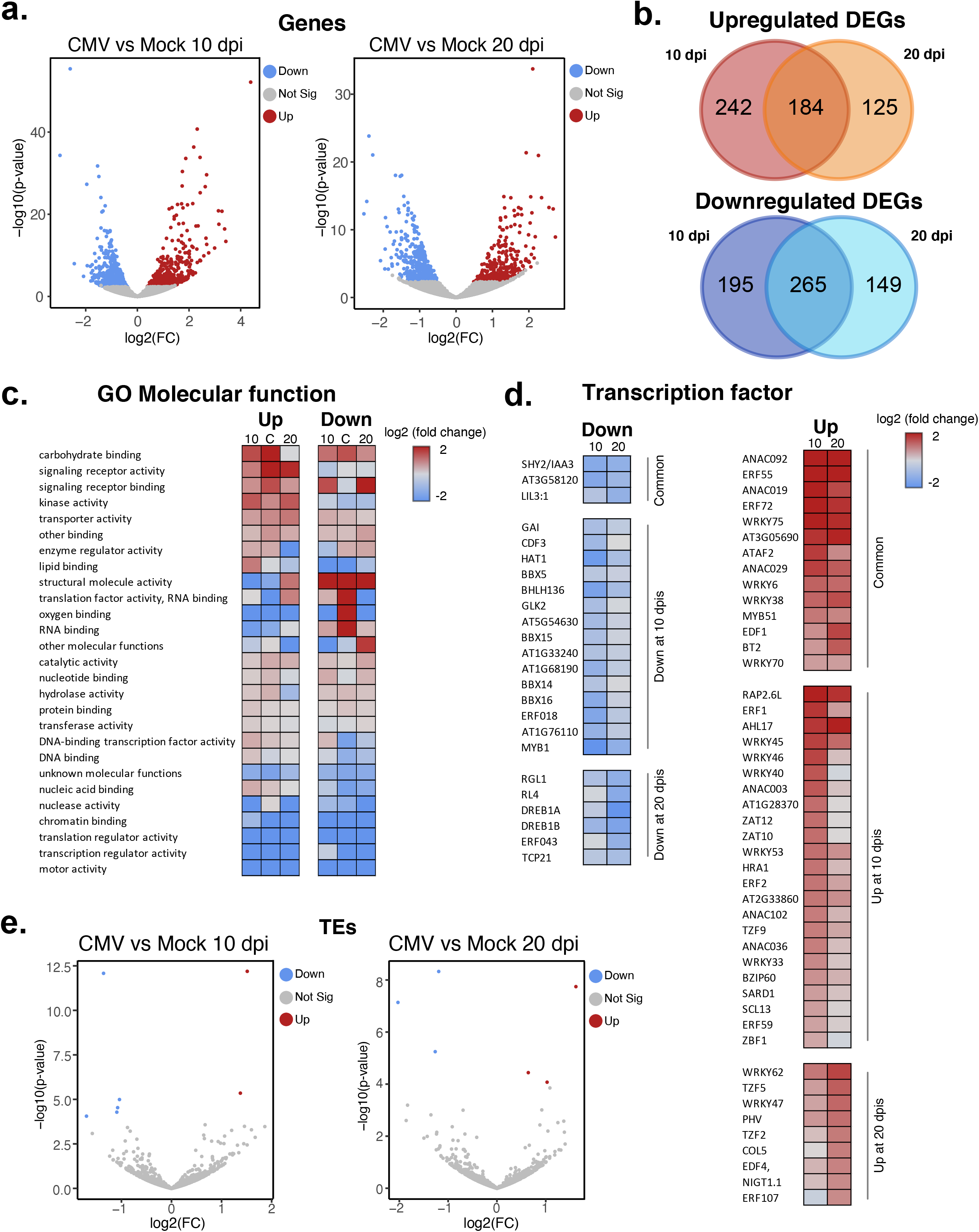
Transcriptomic reprogramming under CMV infection leads to the activation of the defense response. **a.** Volcano plots showing the differentially expressed genes (DEG) at 10 and 20 dpi. **b.** Venn diagrams showing the overall overlap between 10 and 20 dpi upregulated and downregulated DEGs respectively. **c.** Heatmap for the gene ontology (GO) categorization of genes according to their molecular function for DEGs showing upregulation (up) or downregulation (down) at 10, 20 or both times (labeled as 10, 20 or C, respectively). **d.** Heatmaps of the transcription values (showed as log 2 (fold change)) for transcription factors categorized as DEGs showing different patterns of transcriptional activity: downregulation (down) or upregulation (up) and at different time points as indicated. **e.** Volcano plots showing the differentially expressed TEs at 10 and 20 dpi.

Surprisingly, despite causing obvious symptomatology and the misregulation of key epigenetic players, only a modest number of TEs experienced transcriptional changes under CMV infection (Fig 2e). At 10 dpi only 15 TEs were transcriptionally upregulated and 18 TEs were downregulated, while at 20 dpi, 33 TEs were upregulated and 16 TEs were downregulated (Fig 2e and Sup Table 2). Transcriptionally de-regulated TEs at both infection time points were not characterized in a previous study^54^ as targets of the RdDM pathway and instead were classified as TEs with low CHH levels and/or regulated by the CHH-maintenance methyltransferase CMT2 (Sup. Fig 3c). Interestingly, the population of TEs transcriptionally deregulated at 10 dpi showed a less centromeric identity (characterized by shorter length and longer distance to the centromere) compared to the TEs deregulated at 20 dpi, indicating that the mechanisms regulating them might have different dynamics during the viral infection (Sup. Fig 3d). In summary, the results from our transcriptomic analysis reflected a genic transcriptional reprogramming characterized by the elicitation of markers of the defense response against stress and certain deregulation of epigenetic components. In parallel our analysis of TE differential expression pointed to the involvement of multiple epigenetic pathways (other than RdDM) in the control of these repetitive sequences.

### Overall DNA methylation gain during CMV infection

CMV has been previously linked to changes in DNA methylation via its viral silencing suppressor protein 2b^55,56^, which has been proposed to sequester RdDM-derived siRNAs, and as a consequence of the infection *per se*^47,48^. Our WGBS analysis (Fig. 3a) indicated that during CMV infection, at both 10 and 20 dpi, *Arabidopsis thaliana* experienced a significant increase of DNA methylation in all sequence contexts (Fig. 3b and Sup Fig. 4a-b). This increase was more evident at TEs and was higher at 20 dpi, especially in the CHG context (Fig. 3b and Sup Fig. 4b). To further understand the connection between methylation changes and their genomic context, we identified differentially methylated regions (DMRs)^57^. This analysis allowed us to characterize a total of 2768 and 4438 DMRs at 10 and 20 dpi, respectively (Fig. 3c). In accordance with the observed increased levels of DNA methylation, the majority of identified DMRs represent a gain of methylation and are enriched for the CHG context (Fig. 3c and Sup. Fig. 4c). Overall, both gain and loss DMRs in the non-CG contexts are associated with TEs, while gain and loss DMRs in the CG context are mostly associated with genes, especially for the loss of DNA methylation (Fig. 3d). These changes most likely represented the enrichment of each of these sequence contexts among their preferential genomic context, with CG methylation being present at both genes and TEs, while CHG and CHH contexts spreading only through TEs.

**Figure 3.**
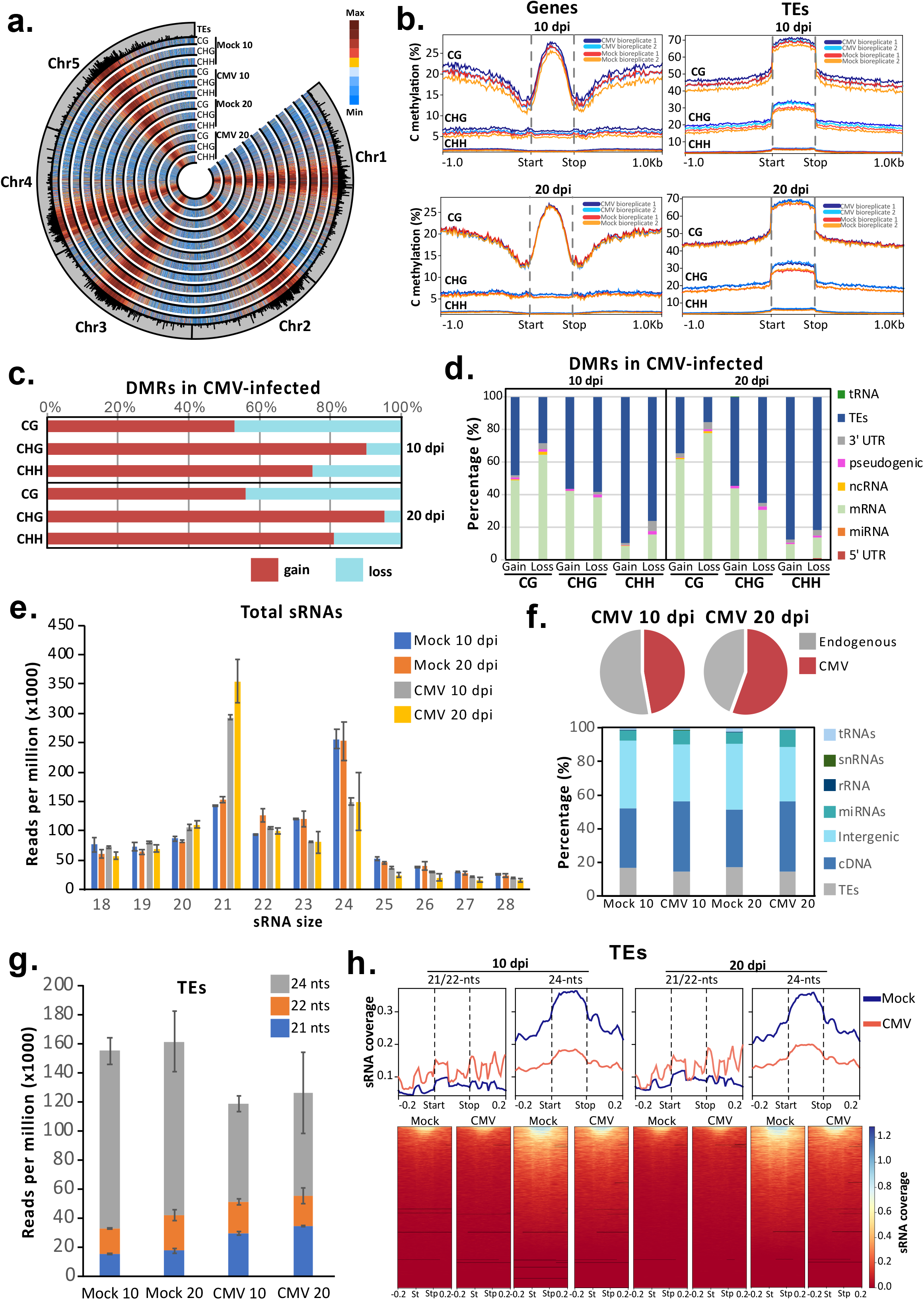
DNA methylation dynamics under CMV infection. **a.** Circular plot showing the genome-wide levels of DNA methylation on each C methylation context on mock and infected samples at 10 and 20 dpi. The outside track shows the localization and genomic length of TEs in the Arabidopsis genome. **b.** DNA methylation coverage profiles of each C methylation context for genes and TEs on mock and infected samples. **c.** Overall presence of gain (hypermethylation) and loss (hypomethylation) differentially methylated regions (DMRs) in CMV-infected samples at 10 and 20 dpi. **d.** Genomic categorization of gain (hypermethylation) and loss (hypomethylation) DMRs identified for each C methylation context on infected samples at 10 and 20 dpi. **e.** Profile of the endogenous sRNA population from 18-to 28-nt in size, normalized to reads per million (x1000), in mock and CMV-infected samples at 10 and 20 dpi. Error bars show the standard deviation between two biological replicates. **f.** Top: pie charts showing the overall presence of endogenous and CMV-derived siRNAs at 10 and 20 dpi; bottom: genomic categorization of the origin of endogenous siRNAs in mock and CMV-infected samples at 10 and 20 dpi. **g.** Distribution of 21-, 22- and 24-nt TE-derived siRNAs in mock and infected samples at 10 and 20 dpi. **h.** Top: sRNA coverage profiles for 21/22- and 24-nt TE-derived siRNAs in mock (blue line) and CMV-infected (red line) samples; bottom: heatmap of the same sRNA coverage profiles represented in the top panel.

DNA methylation is established by siRNAs derived from the RdDM pathway through its canonical and non-canonical forms^58,59^ and maintained by the activity of context-specialized DNA methyltransferases and the same RdDM pathway (at euchromatic loci). To understand the potential connection of siRNAs to the values of DNA methylation observed, we analyzed our high-throughput sRNA data. Our sRNA sequencing (Fig. 3e and Sup. Fig. 5a) indicated striking differences between infected and non-infected tissues at both 10 and 20 dpi. These changes were characterized by a global decrease of 24-nt and an increase of 21-nt endogenous siRNAs (Fig. 3e). A majority of endogenous siRNAs derived from genes and intergenic regions (Fig. 3f and Sup. Fig. 5c), which reflected the changes observed in total siRNAs (Fig 3e). Similar to our previous analysis of CMV virus-derived siRNAs (vsiRNAs)^60^, CMV-infected *Arabidopsis thaliana* plants experienced an overaccumulation of vsiRNAs that increased with the infection time (Fig. 3f and Sup. Fig. 5b).

Changes in sRNA accumulation were connected to loss of DNA methylation since hypomethylated DMRs showed a decrease in 24-nt siRNAs (Sup. Fig. 5d). Interestingly, these changes in 24-nt siRNA accumulation also took place globally over TEs (Fig 3g and h), despite our observed increase in DNA methylation values over these loci (Fig. 3b). Importantly, compared to hypomethylated DMR regions, TEs gained 21-nt siRNAs which could potentially be driving non-canonical RdDM (Fig. 3g and h). In brief, our data indicates that CMV infection led to changes in the global methylation profiles characterized by a gain of DNA methylation at all the different contexts and the presence of abundant hypermethylated DMRs at TEs.

### Dynamic reorganization of H3K9me2 and H3K27me3 during CMV infection

Several components mediating histone homeostasis have been previously involved with the regulation of the stress response in plants. To understand the contribution of histone marks to CMV infection, we focused our analysis on the well-characterized repressive histone marks, H3K9me2 and H3K27me3, which cover the majority of heterochromatin (constitutive and facultative, respectively) in the *Arabidopsis thaliana* genome^61,62^. Our ChIP-seq analysis largely confirmed this trend (Fig. 4a and b). Overall, CMV infection induces an increase of both repressive marks at their targets (Fig. 4b), with an increase of H3K9me2 peaking at 10 dpi and an increase of H3K27me3 at 20 dpi (Fig. 4b and Sup. Fig 6a).

**Figure 4.**
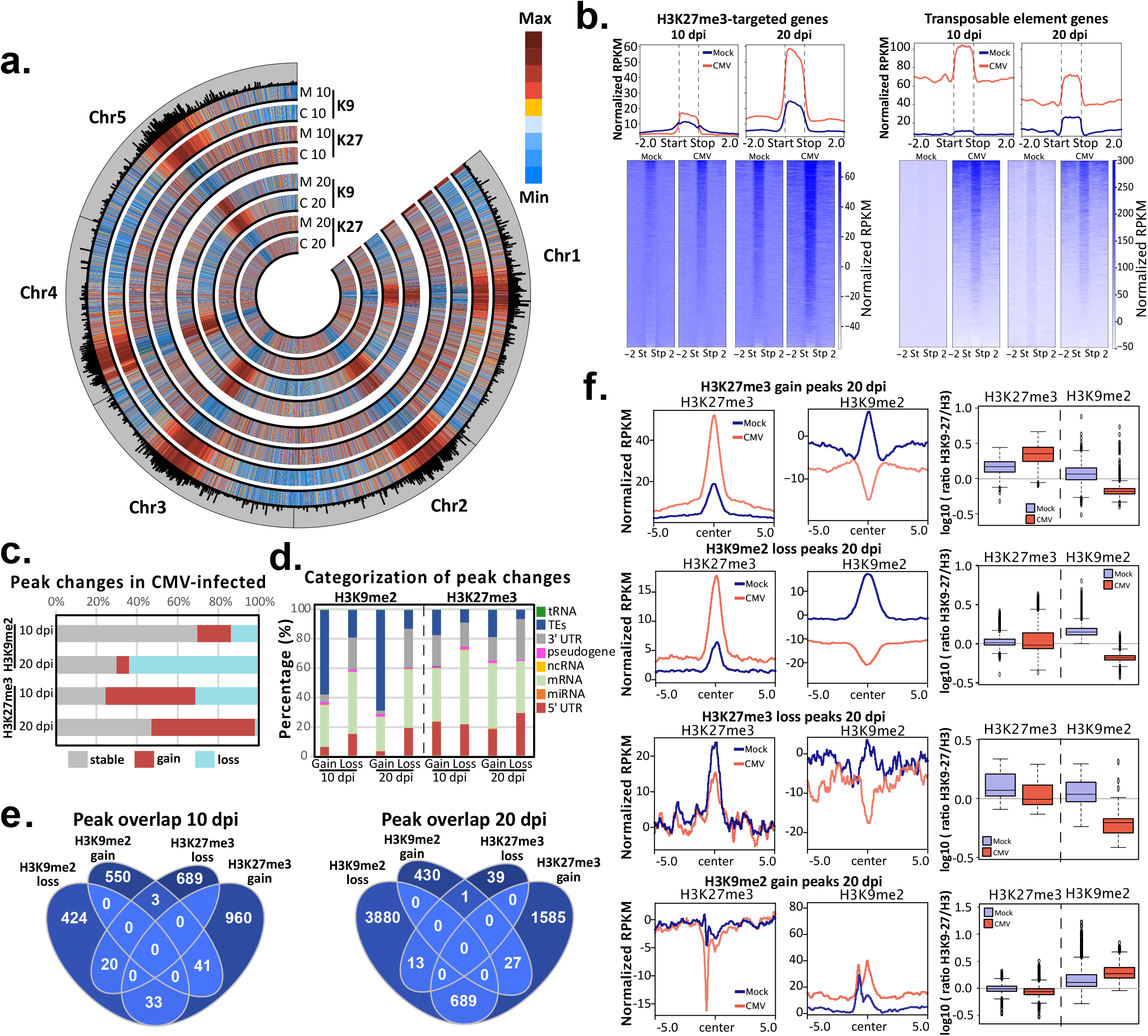
H3K9me2 and H3K27me3 are reorganized under CMV infection. **a.** Circular plot showing the genome-wide levels of H3K9me2 (K9) and H3K27me3 (K27) on mock (M) and CMV-infected (C) plants at 10 and 20 dpi. **b.** Top: histone coverage profiles for H3K27me3-targeted genes (left) and transposable element genes (right) in mock (blue line) and CMV-infected (red line) samples; bottom: heatmap of the same histone coverage profiles represented in the top panel. Values represent RPKM normalized coverage for each mark with the subtracted values of H3 RPKM coverage. **c.** Overall presence of stable, gain and loss H3K27me3 and H3K9me2 peaks in CMV-infected samples at 10 and 20 dpi. **d.** Genomic categorization of gain and loss peaks identified for each histone mark on infected samples at 10 and 20 dpi. **e.** Venn diagram representing the analysis of the overlap of gain and loss H3K9me2 and H3K27me3 peaks at 10 (left) and 20 (right) dpi. **f.** Left panel: H3K27me3 and H3K9me2 coverage profiles at H3K27me3/H3K9me2-gain and loss regions at 20 dpi in mock (blue line) and CMV-infected (red line) samples. Values represent RPKM normalized coverage for each mark with the subtracted values of H3 RPKM coverage; right panel: box plots indicating the overall histone mark values (log 10 of the ratio histone mark/H3-input control). Boxplots are Tukey and p-values were calculated using an unpaired t-test.

To understand in detail the identity of the genomic regions affected by the changes at both histone marks during CMV infection, we identified peaks showing increased and decreased levels of each mark under CMV infection (Fig 4c). While at 10 dpi H3K27me3 showed differences between mock and CMV-infected tissues and H3K9me3 experienced little dynamism, interestingly, at 20 dpi, both histone marks experienced two opposite trends, with a decrease in H3K9me2 peaks and an increase in H3K27me3 (Fig. 4c). In line with their already described preferential association with genomic loci, under viral infection gain of H3K9me2 was mainly located at TEs, while its loss took place at genic regions (Fig. 4d). On the other hand, both gain and loss of H3K27me3 was mostly associated with genes or their regulatory regions (Fig. 4d). Loss of H3K9me2 in mutants that massively lose chromatin or DNA methylation leads to a reorganization of H3K27me3, which invades constitutive heterochromatic regions^63,64^. Our data showed that this interplay between H3K9me2 and H3K27me3 also took place in a wild-type genetic background under CMV infection, since at 20 dpi there was an overlap between regions that lose H3K9me2 and gain H3K27me3 (Fig. 4e and f). This interchange of H3K9me2 by H3K27me3 only took place at regions that had H3K9me2 identity since regions that loss H3K27me3 did not gain H3K9me2 and could retain their facultative heterochromatic identity (Fig. 4f). Gain of H3K9me2 under CMV infection was restricted to very heterochromatic regions of the genome, with already high levels of this histone mark (Fig 4f) and its associated high values of DNA methylation (Sup Fig. 6a). On the other hand, loss of H3K9me2 took place at regions with lower levels of H3K9me2 (Fig. 4f) and relatively low levels of DNA methylation (Sup Fig. 6a). Increased H3K9me2 was associated with a significant increase in DNA methylation in the CHG context at 10 and 20 dpi (Sup Fig. 6a). On the other hand, the loss of H3K9me2 was independent of DNA methylation (Sup Fig. 6a). As expected, H3K27me3 dynamics were completely independent of DNA methylation (Sup Fig. 6b).

In agreement with a loss of heterochromatic identity, reduced levels of H3K9me2 and H3K27me3 lead to higher levels of 21- and 22-nt siRNAs derived from these regions potentially as a result of increased Pol II activity (Sup Fig. 6a and b). Interestingly, high levels of 21- and 22-nt siRNAs were also detected at regions gaining H3K27me3, possibly indicating the relative inability of this repressive mark to restrict Pol II transcription, or its incorporation to reduce previously excessive Pol II transcription under heavy chromatin remodeling. Altogether, our results indicate that histone marks are heavily reorganized during viral infection with stronger changes and interactions between H3K9me2 and H3K27me3 at later infection times.

### DNA methylation and repressive histone marks participate in the transcriptional reprogramming under CMV infection

To understand the influence of DNA methylation, H3K9me2 and H3K27me3 over genic transcription under CMV infection, we analyzed the transcription level of genes that were directly associated with the gain or loss of these repressive epigenetic marks. We focus our analysis on genes that were directly associated with the gain or loss of H3K27me3 within their gene bodies, and genes that gain or loss DNA methylation and/or H3K9me2 within their gene bodies but also within a 1kb window from their gene bodies, since both marks can control regulatory elements influencing gene expression^35^. Following that strategy, we detected 2,192 and 11,356 genes associated with epigenetic marks at 10 and 20 dpi respectively (Sup Fig 7a). The genomic distribution of the genes associated with changes in the epigenetic marks followed the expected location of these marks over the genome, with changes in DNA methylation and H3K9me2 affecting genes significantly closer to the centromere and H3K27me3 affecting genes located further away from centromeres (Fig 5a). Interestingly, following the exchange of repressive histone marks observed at later infection times, at 20 dpi H3K9me2 loss affects genes with slightly less pericentromeric identity and H3K27me3 gain follows the opposite trend (Fig 5a). Several DEGs were found associated with each mark, with a preference for their association with H3K27me3, and different contributions of the other marks (Fig 5b and Sup Table 3). Analysis of the correlation of gene expression with the presence of the different marks showed that indeed H3K27me3 and CG methylation showed the highest significant correlation with their gain or loss and the downregulation or upregulation, respectively, of the corresponding genes (Fig 5c and f). Regarding the rest of the marks, we only observed a correlation between the gain of H3K9me2 and the downregulation of gene expression at 10 dpi, and the gain of CHG methylation and the downregulation of gene expression at 20 dpi (Sup Fig 7c-f). Intriguingly, CHH methylation did not correlate with gene expression changes at any infection time (Sup Fig 7g and h).

**Figure 5.**
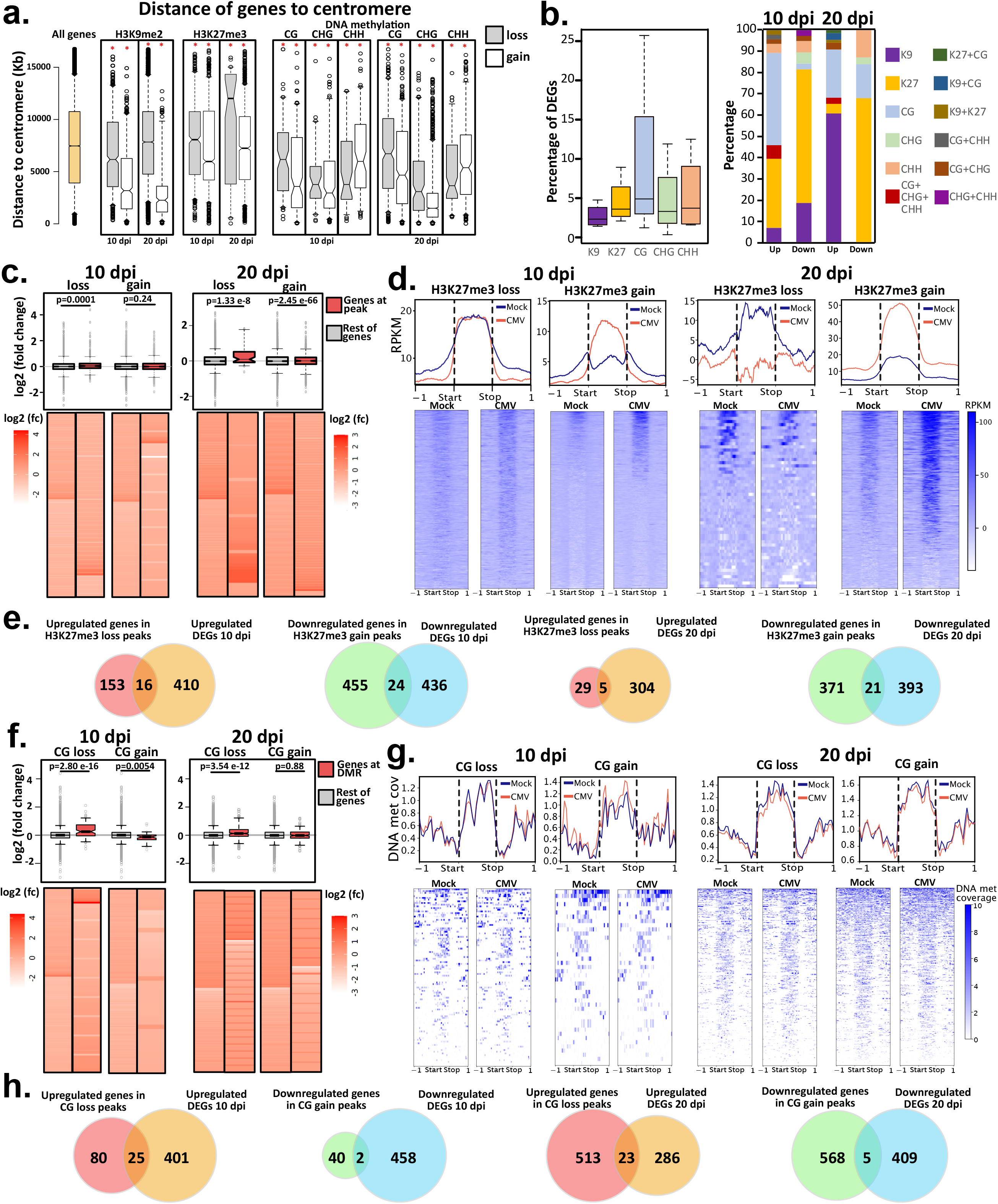
Epigenetic marks contribute to the transcriptional reprogramming under CMV infection. **a.** Box-plots depicting the overall distance to the centromere for all genes associated with each of the marks at the infection times indicated. The asterisk indicates p-value<0.05 calculated through an unpaired t-test comparing the distance for each mark against the distance for all genes in the *Arabidopsis thaliana* genome (yellow box). **b.** Left: Box-plots indicating the percentage of DEGs within the overall genes associated with each epigenetic mark indicated in the color legend. Right: Association of the DEGs with each epigenetic mark under study here as indicated in the color legend. **c.** Expression values (log 2 (fold change)) for all genes associated with H3K27me3 loss and gain peaks at 10 and 20 dpi (colored in red) or the rest of the genes (all genes in the *A.thaliana* genome with the subtracted values for peak-associated genes). p-values are indicated on top of each comparison and were calculated through an unpaired t-test. **d.** Top: histone coverage profiles for genes associated with a gain or loss H3K27me3 peak at 10 and 20 dpi in mock (blue line) and CMV-infected (red line) samples; bottom: heatmap of the same histone coverage profiles represented in the top panel. Values represent RPKM normalized coverage for each mark with the subtracted values of H3 RPKM coverage. **e.** Venn diagrams depicting the overlap between genes showing the expected upregulation or downregulation upon H3K27me3 loss or gain, respectively and upregulated or downregulated DEGs at the respective infection times indicated. **f.** Expression values (log 2 (fold change)) for all genes associated with CG loss and gain peaks at 10 and 20 dpi (colored in red) or the rest of the genes (all genes in the *A.thaliana* genome with the subtracted values for DMR-associated genes). p-values are indicated on top of each comparison and were calculated through an unpaired t-test. **g.** Top: DNA methylation coverage profiles for genes associated with a CG gain or loss DMR at 10 and 20 dpi in mock (blue line) and CMV-infected (red line) samples; bottom: heatmap of the same DNA methylation coverage profiles represented in the top panel. Values represent DNA methylation coverage. **h.** Venn diagrams depicting the overlap between genes showing the expected upregulation or downregulation upon CG loss or gain, respectively and upregulated or downregulated DEGs at the respective infection times indicated.

DEGs regulated by epigenetic marks were confirmed to be targets of their respective pathways since they were identified as differentially expressed (adjusted p-value <0.05) in different combinations of mutants for genes involved in the homeostasis of DNA methylation (*mddcc*^64^, *ago4*^65^, and *polV*^66^, 71% of DNA methylation-associated DEGs), H3K27me3 (*clf* and *elf6 ref6 jmj13*^67^, 93% of H3K27me3-associated DEGs) and H3K9me2 (*kyp* and *ddm1*^68^, 54% of H3K9me2-associated DEGs) (Fig 6a and Sup Table 4). The majority of overrepresented CMV-induced DEGs associated with epigenetic marks were classified as stress/stimulus responsive genes according to their annotated gene ontology (GO) biological function (Fig 6b and Sup Fig 8a). Interestingly, certain groups of genes were predominantly associated with some epigenetic marks, such as secondary metabolic processes (H3K27me3-associated) or cell death (DNA methylation- and H3K9me2-associated) (Fig 6b). A similar result was obtained when genes were grouped by molecular function (Sup Fig 8b). Reflecting our identification strategy, genes regulated by DNA methylation and H3K9me2 included genes with increased/decreased gene body and TE-like DNA methylation/H3K9me2 of elements surrounding their regulatory regions (Sup Fig 8c-d). In comparison H3K27me3 was only found associated with its presence in the gene body (Sup Fig 8e). DEGs regulated by epigenetic marks included several important genes for the biology of viral infection such as the previously identified virus-responsive genes WRKY40^69^ (associated with DNA methylation, Fig 6c); WAK1^70^, NHL10^71^, CHI^72^ (all associated with H3K27me3, Fig 6d); and PUM5^73^, HSP70-2^74^ and the main antiviral AGO protein, AGO2^75^ (all associated with H3K9me2, Fig 6e). In addition, CMV-responsive DEGs associated with epigenetic marks included other stress-responsive genes that have not been previously associated with viral infections such as AT1G43910, AT1G13470 and AT1G67870, the lipid transfer protein EARLI1, the acyl-transferase protein CER26 or the FAD-binding Berberine family protein ATBBE10 (Sup Fig 8c-e).

**Figure 6.**
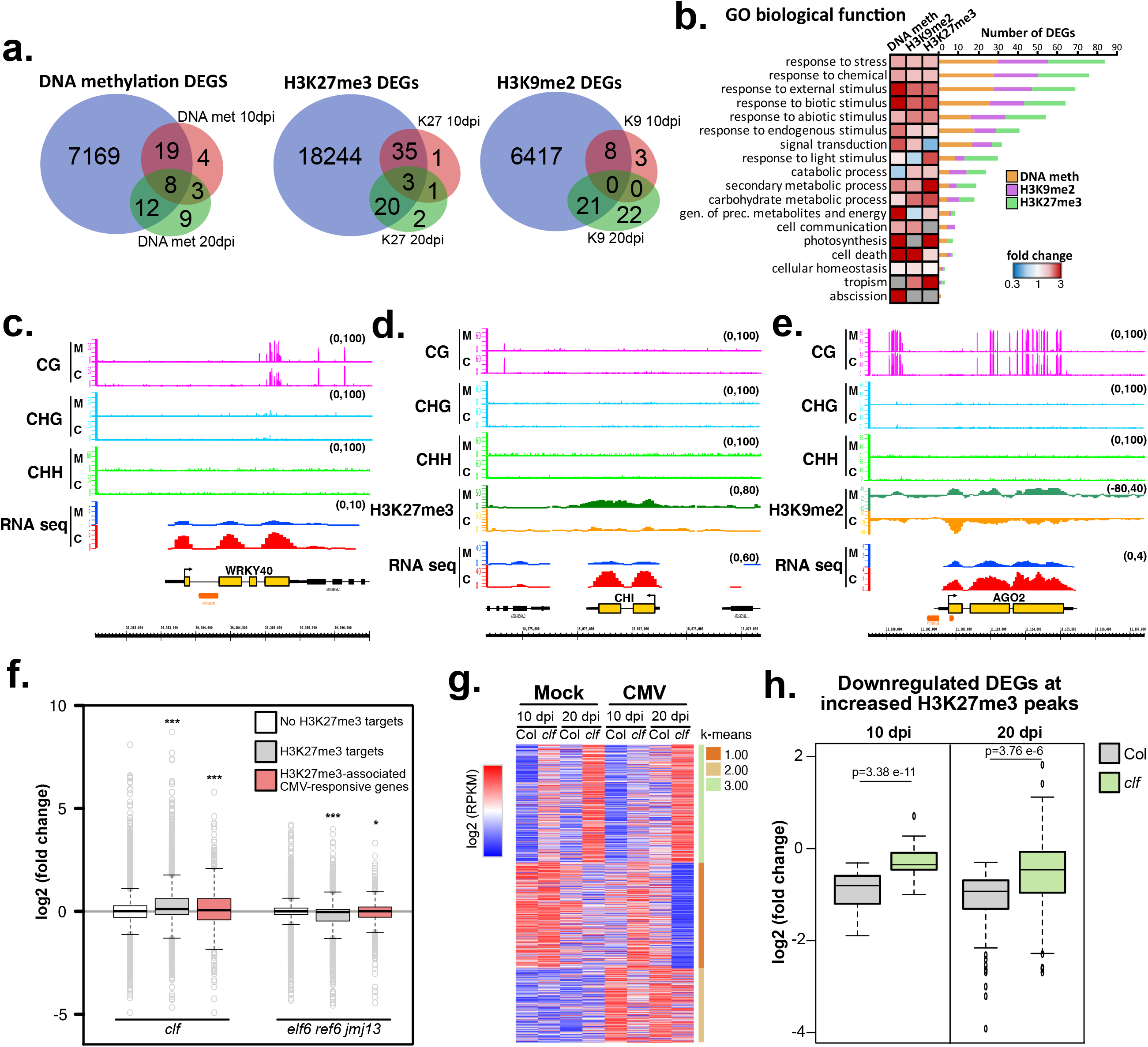
Epigenetic marks regulate multiple differentially expressed genes through different mechanisms and CLF is mechanistically involved in downregulating H3K27me3-associated genes during CMV infection. **a.** Venn diagrams showing the overlap between DEGs associated with different epigenetic mechanisms and CMV-induced DEGs associated with the different epigenetic marks under study here (DNA methylation, H3K27me3 and H3K9me2). **b.** Heatmap for the gene ontology (GO) categorization of genes according to their biological function for DEGs associated with DNA methylation, H3K27me3 or H3K9me2. Number of genes associated with each category is shown on the right of the heatmap, with the colored number of genes associated with each specific mark. **c-e.** Genome browser screenshots showing the association of different viral-responsive DEGs previously identified as important in the viral infection process and associated with decreased values of DNA methylation in the CG context at 10 dpi for WRKY40 (c), H3K27me3 at 20 dpi for CHI (d) and H3K9me2 at 20 dpi for AGO2 (e). M=mock and C=CMV. **f.** Box-plots showing expression values (log 2 (fold change)) for genes identified as non-H3K27me3 targets (white boxes), H3K27me3 targets (grey boxes) or H3K27me3 targets during CMV infection (red boxes) in *clf* and *elf6 ref6 and jmj13* mutant backgrounds. Asterisks show different p-values: *<0.05, ***<0.005. **g.** Heatmap showing the RNA expression values (log 2 of the RPKM values) for all H3K27me3-associated genes (as described in Zhang et al 2007^62^) under CMV-infection in Col or *clf* genetic backgrounds in mock or CMV-infected conditions at 10 and 20 dpi as indicated. **h.** Box-plots showing expression values (log 2 (fold change)) of DEGs downregulated by increased values of H3K27me3 under CMV infection in Col (grey boxes) and *clf* (green boxes) genetic background. p-values are indicated on top of each comparison and were calculated using an unpaired t-test.

These results highlight the potential of epigenetic changes mediated by multiple epigenetic marks in regulating the transcriptional activity under CMV infection. In particular, we found that H3K27me3 and CG methylation changes were highly correlated with the expression of their associated genes.

### CLF represses gene expression during CMV infection

Next, we aimed to understand the mechanistic contribution of CLF (identified in our initial screening for mutants showing resistance to CMV infection, Fig 1a and b) to the resistance to CMV infection. H3K27me3 profiles under CMV infection showed a pattern compatible with an important role of this mark during CMV infection (Fig 4 and 5). Indeed, H3K27me3-associated genes during CMV infection were bona-fide H3K27me3 targets since they behave similarly to H3K27me3 targeted genes and were upregulated in a *clf* mutant background but downregulated in a triple *elf6 ref6 jmj13* mutant (Fig 6f). To understand the role of CLF during CMV infection, we infected *clf* mutants with CMV and produced high-throughput RNA sequencing libraries at 10 and 20 dpi (Fig 6g and Sup Fig. 9a-e). Our analysis of gene expression indicated that *clf* mutants showed increased expression level of H3K27me3 targets (Fig 6g and Sup Table 5), which is significantly higher for DEGs that were downregulated in associated with increased H3K27me3 at both 10 and 20 dpi (Fig 6h). In sum, our data shows that CLF is an important regulator of the transcriptional reprogramming taking place under CMV infection and that it negatively regulates the expression of genes targeted by H3K27me3 during infection.

## Discussion

Epigenetic mechanisms regulate important aspects of cellular viability such as the genome stability and/or accessibility to the transcriptional machinery^76,77^. Furthermore, epigenetic mechanisms are dynamic and provide a layer of versatility to rapidly adapt to environmental signals, including stresses^19–21,78^. Indeed, the role of DNA methylation in regulating gene expression under different biotic and abiotic stresses has been extensively described, while the role of other epigenetic marks has remained uncharacterized. Here we have used CMV infection progression (a well-studied biotic stress) to analyze the genome-wide changes in multiple epigenetic marks and their connection to transcriptional activity (using both RNA and sRNA sequencing).

We find that under CMV infection, the repressive histone marks H3K9me2 and H3K27me3 are more dynamic than DNA methylation, and that H3K27me3 plays a role in the resistance to CMV infection. In line with this, we find that CLF (a catalytic component of the PRC2 complex) controls the downregulation of genes silenced by the increase of H3K27me3 during infection. Our data also exemplifies the dynamism of epigenetic marks, which (in a wild type genetic background) are extensively and functionally reorganized.

Our analysis also adds more information about the particular role of epigenetic marks during stress. In our pathosystem, we did not identify major changes in DNA methylation, with only an overall increase in DNA methylation observed under CMV infection (Fig 3). These changes, which occur mostly in the CHG context might be attributable to our observed reorganization of H3K9me2 during CMV infection (Fig 4) rather than to differential activity of the RdDM pathway. In line with this observation, most of the TEs differentially expressed under viral infection are not regulated by the RdDM pathway and they either, have low CHH methylation levels or their methylation is maintained independently of the RdDM pathway by CMT2. The lack of more widespread TE transcriptional reactivation during CMV infections is surprising due to the strong decrease of 24-nt TE-derived siRNAs observed (Fig 3e). We find plausible that TE transcriptional silencing under CMV infection could be a consequence of this strong H3K9me2 increase in centromeric and pericentromeric regions at both 10 and 20 dpi. Alternatively, DNA methylation increase could be attributable to either the action of a non-canonical version of the RdDM pathway using 21-/22-nt siRNAs to introduce DNA methylation or an overall hyperactivity of the maintenance methylation pathways. Previous works have shown that CMV infection dramatically alters the sRNA landscape of the infected cells, leading to up to 50% of the cellular sRNAs deriving from the viral genomic RNAs^60^. These dramatic alterations of the overall siRNA profiles lead to the leaking of vsiRNAs into almost all endogenous AGO proteins, potentially affecting (directly and indirectly) multiple cellular processes^60^. Additionally, CMV viral silencing suppressor, 2b, is a known interactor of the PTGS machinery, sequestering sRNAs of different sizes and classes, including endogenous 24-nt siRNAs^60^. We discard that Dicer-independent DNA methylation might occur in our tissues, since no laddering of endogenous siRNAs (characteristic of this type of DNA methylation pathway) was observed in our sRNA sequencing data. Our data also highlights the need to consider the possibility that the dynamism in DNA methylation induced by multiple stresses could, at least partially, be the result of genome-wide histone reorganization.

Furthermore, our data identified a compensatory mechanism between H3K9me2 and H3K27me3 (Figure 4e-f). This indicates that compensation of these heterochromatic marks is functional and takes place in a wild-type genetic background (under CMV infection) and is not only an artifact occurring in mutant backgrounds that have compromised chromatin such as *mddcc*^64^ or *ddm1*^63^. A similar functional compensatory mechanism between H3K9me2/3 and H3K27me3 has previously been observed in the silencing of the chromosome X, which shows both high levels of H3K9me3 and H3K27me3 and a complementation of both marks to mediate the X inactivation process^79^.

Finally, we found that the changes in all the repressive marks analyzed were connected to the transcriptional response during viral infection and we found examples of genes regulated by both DNA methylation and repressive histone marks (Fig 5). During the progression of CMV infection, both DNA methylation and H3K9me2 gradually associate with genes that have a stronger pericentromeric and centromeric location. At 20 dpi, H3K27me3 gain also leaks into pericentromeric locations, probably to compensate the re-organization of H3K9me2 (Fig 5a). Indeed, several DEGs are regulated by H3K27me3 (Fig 6), the mark that better correlated with the direction of gene expression (Fig 5c), which is expected since this epigenetic mark is key to regulate transcriptional networks^80^. DEGs regulated by H3K27me3 include the upregulated chitinase CHI (Fig 6d), which is responsive to different viruses^72^; the upregulated gene ATBBE10, a FAD-binding Berberine protein member that is responsive to multiple biotic and abiotic stresses^81–83^, epigenetically regulated^81^ and whose overexpression is associated with resistance to bacteria^84^; and the downregulated gene CER26 a HXXXD-type acyl-transferase protein^85^ that is positively regulated by histone acetylation and involved in stem cuticular wax biosynthesis^86^. Interestingly, we found that the re-organization of

H3K9me2 is associated with the overexpression of AGO2, the main antiviral AGO protein, which is indeed a gene with pericentromeric location that contains multiple TEs in its 5’ regulatory region, a fingerprint of epigenetically-regulated genes (Fig 6e and Sup Fig 8f). We found that downregulated DEGs under CMV infection (such as CER26) are targeted by CLF, indicating that this catalytic member of PRC2 is needed in the orchestration of the transcriptional reprogramming under viral infection and explaining the observed increased resistance of CLF mutants to CMV. Rewiring of H3K27me3 and H3K9me2 could also be partially responsible for the transgenerational memory of stress that has been observed in plants exposed to different stresses^87–92^, since both these marks can be partially inherited to the next generation^93,94^.

Based on our results, we speculate that the hijacking of the cellular machinery by CMV leads to different simultaneous epigenomic consequences (Sup Fig 10). First, the previously characterized interference of CMV with the RdDM machinery caused by its viral suppressor of RNA silencing, the 2b protein, or the generalized loading of vsiRNAs (and displacement of the endogenous siRNAs) into the endogenous AGO proteins, might lead to reduced DNA methylation at epigenetically labile loci. These changes in DNA methylation might lead to a preventive re-organization of H3K9me2 intro pericentromeric and centromeric regions to avoid the transcriptional activation of TEs. This reorganization of constitutive chromatin might lead to two consequences for genic activity: a) a transcriptional activation of pericentromeric genes, and b) a compensatory activity of H3K27me3 to silence excessive pericentromeric gene activity mediated (at least partially) by CLF. Our work exemplifies that epigenetic changes associated with the stress response are not limited to DNA methylation and that the interplay between epigenetic marks during stress is extensive. Furthermore, our results provide a step further into understanding the interaction between plants and viruses and the dynamism of the epigenome during stress.

## Methods

### Plant material, CMV infection and phenotypic measurements

*Arabidopsis thaliana* (Columbia wild-type Col-0) were sown into potting soil (P-Jord, Hasselfors Garden, Örebro, Sweden). At the four-leaf stage, seedlings were selected by uniformity and carefully replanted into plastic pots (9 × 9 × 7 cm) with one plant per pot at temperature 20–22°C and 45% relative humidity. Plants were grown under a 16 h : 8 h, light : dark photoperiod. The light was provided by FQ, 80 W, Hoconstant lumix (Osram, Munich, Germany) with a light intensity of 220 μmol photons m^−2^ s^−1^. Mutants alleles used in this work include *drm1-2 drm2-2 cmt3-11*^95^ (*ddc*), *ddm1-2*^96^, *kyp-6*^97^ (Salk_041474), *nrpd1a-3*^98^ (Salk_128428), *clf-29*^99^ (Salk_021003), *ref6-1*^100^ (Salk_001018), *ago4-5*^101^ (CS9927), *cmt3-11*^102^ (CS16392) *and drm2-2*^103^ (Salk_150863). CMV infection was performed in plants at the 4 rosettes leaves stage (1.04 stage from Boyes et al. 2001^104^). Two leaves per *Arabidopsis thaliana* (Columbia wild-type Col-0) were rub-inoculated with a sap-solution obtained by homogenizing previously infected *Nicotiana benthamiana* leaves on 0.1 M Na_2_HPO_4_. Mock plants were rub-inoculated only with the 0.1 M Na_2_HPO_4_ buffer. *N. benthamiana* plants were infected with the three genomic RNAs of CMV strain FNY, previously obtained by *in vitro* transcription from plasmids containing the individual genomic sequences using the MAXIscript T7 Transcription Kit (Thermo Fisher). *N. benthamiana* symptomatic leaves were collected at 12 dpi and kept at -70 °C as a viral reservoir. *A. thaliana* samples were collected at 10 and 20 dpi. To determine the degree of the infection at a phenotypic level, measurements of the radius of the rosettes were taken at both 10 and 20 dpi. For that, two measurements for the rosette were taken, trying to take them in a cross shape as much as possible. The final measurements were calculated by doing the mean of the two numbers. Statistical significance was calculated using unpaired *t*-tests.

### DNA/RNA extraction and RT-qPCR

Genomic DNA for regular genotyping and bisulfite sequencing was extracted using the DNeasy Plant Mini Kit (Qiagen) following the manufacturer’s instructions. Total RNA was extracted using TRIzol reagent (Life Technologies) following the manufacturer’s instructions. For RNA sequencing, mRNA was purified with the NEB mRNA isolation kit (New England Biolabs). Extracted RNA was treated with DNase I (Thermo Fisher) and used to synthesize cDNA using the RevertAid First Strand cDNA Synthesis Kit (Thermo Fisher), following the manufacturer’s instructions. Then, cDNA levels were measured using the 5x FIREPol EvaGreen qPCR Mix Plus (ROX) (Solis Biodyne). Finally, relative accumulation was calculated using the “delta-delta method” formula (2^-<[^ ^ΔCP^ ^sample-ΔCP^ ^control]^), where 2 represents a perfect PCR efficiency. *UBQ10* (primers used AAGCAGTTGGAGGATGGCAGAAC (forward) and CGGAGCCTGAGAACAAGATGAAGG (reverse)) was used as the housekeeping gene to which the levels of viral cDNA (primers used CTTCCAGAGATGCCTTCGAG (forward) and GGCAGTGCTTGTTCTTGACA (reverse)) were normalized. Statistical significance was calculated using unpaired *t*-tests. For each mutant, three biological replicates consisting of a pool of 8-10 plants and three technical replicates were used.

### Small RNA and RNA libraries preparation, sequencing, and analysis

Small RNA libraries were prepared using the NEBNext® Small RNA Library Prep Set for Illumina® (New England Biolabs), and each individual library was barcoded using the NEBNext® Multiplex Oligos for Illumina® kit (New England Biolabs). RNA libraries were prepared using the NEBNext® Ultra™ II Directional RNA Library Prep Kit for Illumina® (New England Biolabs) and each individual library was barcoded using the NEBNext® Multiplex Oligos for Illumina® kit (New England Biolabs). For both types of libraries, two biological replicates consisting of pools of 3-4 plants per condition were sequenced. The obtained sequences were pre-processed by the following bioinformatics workflow: de-multiplexed, adapter trimmed, and filtered by length and quality. Afterwards, the remaining sequences were aligned to the *Arabidopsis* genome. For sRNA seq analysis, the alignment was performed using bowtie^105^ with the following parameters: -t -v2, allowing two mismatches. Library size was normalized by calculating reads per million of 18-28 nt genome-matching sRNAs.

RNA sequencing libraries were sequence as paired-end 150 bp fragments in an Illumina Novaseq 6000at Novogene (Beijing, China). The obtained raw reads were trimmed using Trimgalore 0.6.1 for the removal of the adapter sequences. For expression analysis, paired reads were aligned to the Arabidopsis TAIR10 genome using STAR^106^ with the following parameters: --outMultimapperOrder Random --outSAMmultNmax -1 -- outFilterMultimapNmax 100 or default(10) -outSAMattributes NH HI AS NM MD -- outSAMunmapped Within --outSAMtype BAM SortedByCoordinate --quantMode TranscriptomeSAM GeneCounts --outWigType bedGraph --limitBAMsortRAM 24000000000. The parameter -N10 or -N100 were used for the analysis of gene or TE expression, respectively. Afterwards, count reads per gene were obtained using HTSeq-COUNTS^107^ with the following parameters: --mode union --stranded no --minequal 10 and --nonunique none. For TE expression analysis, paired reads were aligned to the Arabidopsis TAIR10 genome using STAR, allowing the mapping to at most 100 ‘best’ matching loci with the following parameters: --outMultimapperOrder Random --outSAMmultNmax -1 -- outFilterMultimapNmax 100, as previously used in Warman *et al.* 2020 ^108^. The count reads per TE were obtained using HTSeq-COUNTS using the following parameters: --mode union --stranded no --minequal 0 and --nonunique all. The obtained count tables were used in DESeq2^109^ to infer significant expression with fit type set to parametric. All these tools were used on the Galaxy platform^110^. Volcano plots were created using the R package ggplot2^111^.

### Bisulfite library preparation and sequencing analysis

Bisulfite libraries of two bioreplicates per condition were produced from genomic DNA and sequence as paired-end 150 bp fragments in an Illumina Novaseq 6000at Novogene (Beijing, China). The obtained raw reads were trimmed using Trimgalore 0.6.1 for the removal of the adapter sequences and 10 bases from 5’ ends. The remaining sequences were aligned to the Arabidopsis TAIR10 genome using Bismark^112^, allowing one mismatch per 25 nt seed and keeping the forward and reverse reads independently mapped. Cytosine conversion rates were obtained using bismark_methylation_extractor; the first seven bases from the 50 end and 13 from the 30 end of each read were ignored. The mean conversion rate based on the cytosine methylation levels in the chloroplast genome for the four samples was 99.76%, and the estimated false-positive methylation rates were 0.24%.

For DMR identification, the genome was divided into equal bins of 50 bp in size and the biological replicates from each condition were pooled and compared. Then, the DMRs were identified by performing Fisher’s exact test between the number of methylated reads and the total number of reads in both conditions for each bin. The obtained p-values were adjusted for multiple testing using Benjamini and Hochberg’s method to control the false discovery rate ^113^. Bins with fewer than three cytosines in the specified context or < 0.25 difference in methylation proportion between the two conditions or an average number of reads lower than 8 were discarded. Finally, bins that were at <300 bp were joined.

### Chromatin Immunoprecipitation (ChIP) sequencing libraries preparation and sequence analysis

500 mg of rosette leaves were chemically cross-linked using 1% formaldehyde. Nuclei were isolated from cross-linked material following a standard nuclei isolation protocol based on sucrose gradients as previously described^114^. Resuspended nuclei pellets were sonicated for 9 cycles of 20 s On and 45 s Off at 4 °C and high. Afterward, the Immunoprecipitation (IP) was performed following a standard IP protocol and using the following antibodies: H3 (Reference: 07-690, Merck), H3K9me2 (Reference: pAb-060-050, Diagenode) and H3K27me3 (Reference: 07-449, Merck). The resulting immunocomplexes were purified with the GeneJET PCR Purification Kit (Thermo Fisher), following the manufacturer’s instructions. Finally, DNA libraries were prepared using the NEBNext® Ultra™ II DNA Library Prep Kit for Illumina® (New England Biolabs) and each individual library was barcoded using the NEBNext® Multiplex Oligos for Illumina® kit (New England Biolabs).

ChIP libraries of two bioreplicates per condition were obtained from the immunoprecipitated DNA and sequenced as paired-end 150 bp fragments in an Illumina Novaseq 6000 at Novogene (Beijing, China). The obtained raw reads were trimmed using Trimgalore 0.6.1 to remove the adapter sequences and 10 bases from 5’ ends. For genome-wide distribution analysis sequences were aligned to the Arabidopsis TAIR10 genome using bowtie2 with default parameters. BAM files were filtered for unique reads using the parameters -q20 -F255 -f0X2 and replicates were merged using samtools^115^. Genome coverage was calculated as the log2 fold change of the ratio between the coverage of H3K9me2 or H3K27me3 to the coverage of H3 using deeptools2^116^. For genome-wide profile images, the RPKM-normalized value of H3 was subtracted to the RPKM value of either H3K9me2 or H3K27me3. Values for specific regions and quantitative analysis were retrieved using mapbed from bedtools ^117^.

Peak calling was performed using Sicer2 each sample to its respective H3 control with the parameters window size 200, fragment size 150, effective genome fraction 0.74, false discovery rate 0.01, false discovery rate of 0.01 and a gap size of 600bp. Peak location and overlap was compared using the intersect tool from bedtools with a minimum overlap of 1bp. Only peaks shared between the two replicates were considered as true peaks for that specific treatment. Shared peaks were compared between samples using the intersect tool from bedtools to determine gain and loss peaks.

### Gene ontology (GO) term analysis

GO term analysis was performed using the *GO annotation search, functional categorization and download* tool from the TAIR website (www.arabidopsis.org). Bar plots were created using the R package ggplot2 ^111^.

### Cytology

Immunostaining of *Arabidopsis thaliana* nuclei was performed as previously described^118^. In brief, 6-week old (mock and CMV-infected) rosette leaves were collected and fixed in cold 4% paraformaldehyde in Tris-HCl buffer (10mM Tris pH 7.5, 10 mM EDTA and 100 mM NaCl) for 20 minutes followed by two washes with ice-cold Tris-HCl buffer twice for 10 minutes each. Nuclei were isolated by chopping leaves in LB01 buffer (15mMTris-HCl pH7.5, 20mM NaCl, 2mM EDTA, 80mM KCl, 0.5mM spermine, 0.1% TritonX-100) and filtered through a 30-μm Cell Trics filter (Sysmex, Germany). The filtered nuclei were diluted 1:3 with sorting buffer (100mM Tris pH7.5, 50mM KCl, 2mM MgCl2 2, 0.05% Tween 20, 5% sucrose), and spotted onto microscopy slides to air-dry. The slides were post-fixated with 4% paraformaldehyde for 15 min at room temperature in PBS buffer (10 mM sodium phosphate, pH 7.0, 143 mM NaCl), and washed twice with PBS for 5 minutes each. Slides were blocked with 4% BSA for 30 minutes at 37°C in a moist box followed by primary (anti-H3K9m2, C15410060, Diagenode, 1:500) and secondary (Alexa Fluor 488 Goat anti-Rabbit IgG1 Secondary Antibody, A21121, Invitrogen, 1:100) antibody incubation (all diluted in 1% BSA, 0.1% tween20, 1x PBS). After the antibody incubations, slides were mounted with 2 μg/ml DAPI and analyzed in a confocal microscope (Zeiss LSM780).

## Acknowledgements

We thank Formas (2021-01161), the Swedish Research Council (VR 2021-05023), and the Knut and Alice Wallenberg Foundation (KAW 2019.0062) for supporting research in the Martinez group. Sequencing was performed at Novogene (China). Open Access funding provided by Swedish University of Agricultural Sciences. The data handling was enabled by resources provided by the Swedish National Infrastructure for Computing (SNIC) at UPPMAX partially funded by the Swedish Research Council through grant agreement no. 2018-05973. The authors want to dedicate this manuscript to the loving memory of Juan Luis Reig-Valiente.

